# Accurate identification of medulloblastoma subtypes from diverse data sources with severe batch effects by RaMBat

**DOI:** 10.1101/2025.02.24.640010

**Authors:** Mengtao Sun, Jieqiong Wang, Shibiao Wan

**Affiliations:** Department of Genetics, Cell Biology and Anatomy, University of Nebraska Medical Center, Omaha, NE, USA; Department of Neurological Sciences, University of Nebraska Medical Center, Omaha, NE, USA

**Keywords:** Medulloblastoma, Batch effect, Intra-sample gene ranking, Cancer subtype identification, RaMBat

## Abstract

As the most common pediatric brain malignancy, medulloblastoma (MB) includes multiple distinct molecular subtypes characterized by clinical heterogeneity and genetic alterations. Accurate identification of MB subtypes is essential for downstream risk stratification and tailored therapeutic design. Existing MB subtyping approaches perform poorly due to limited cohorts and severe batch effects when integrating various MB data sources. To address these concerns, we propose a novel approach called RaMBat for accurate MB subtyping from diverse data sources with severe batch effects. Benchmarking tests based on 13 datasets with severe batch effects suggested that RaMBat achieved a median accuracy of 99%, significantly outperforming state-of-the-art MB subtyping approaches and conventional machine learning classifiers. RaMBat could efficiently deal with the batch effects and clearly separate subtypes of MB samples from diverse data sources. We believe RaMBat will bring direct positive impacts on downstream MB risk stratification and tailored treatment design.

## Introduction

As the most common malignant pediatric cancer arising in the cerebellar vermis, medulloblastoma (MB) accounts for around 20% of all pediatric central nervous system (CNS) neoplasms and over 70% of patients diagnosed are children under the age of 10 ^1,2^. Based on the genetic heterogeneity and phenotypic diversity ^3^, four major MB subgroups with distinctive molecular landscapes, somatic mutations and clinical outcomes are defined by the World Health Organization (WHO) ^4^, including Sonic HedgeHog (SHH) activated, Wingless-Type (WNT) activated, and the numerically designated Group 3 and Group 4 ^5^. WNT and SHH are named after signaling pathway disturbances in the WNT (*CTNNB1* mutation ^6,7^) and SHH (*PTCH*, *SUFU*, *SMO* mutation, or *GLI2* amplification ^8^) respectively, which are clearly separable with other two subtypes. Currently, the long-term survival rate of MB has reached 70% based on the standard treatment, however, there are wide disparities among patient outcomes due to the significant difference in histologic characteristics of the tumor, age, residual disease and other factors ^9,10^. WNT can be almost completely cured under current therapy schemes (5-year event-free survival greater than 90% ^11^), whereas SHH-activated MB occurs more frequently in infants and largely dependent on the specific genetic features where *TP53* mutation caused tumor has poorer prognosis ^8^. Group 3 and Group 4 do not exhibit subgroup-defining mutations ^12^, but still have distinct clinic-biological features where Group 3 patients have the most unfavorable prognosis due to a higher incidence of “high-risk” features, such as LCA (large cell/anaplastic) histology and *MYC* amplification ^5,13–15^, and Group 4 has an intermediate prognosis with the feature of isochromosome 17q (*i17q*) ^16^. In other words, the survival rates vary significantly with different MB subtypes. Conventional radiation and chemotherapy have been demonstrated to have long-term sequelae. Given the heterogeneity of these molecular subtypes, tailoring individualized treatment plans not only enhances therapeutic efficacy but also minimizes the long-term side effects associated with conventional therapies ^17^. Therefore, accurately identifying MB subtypes is crucial for guiding effective treatment strategies.

Conventional methods like morphological analysis, immunophenotyping, or molecular profiling for identifying MB subtypes are costly, time-consuming, or inaccurate. To overcome these problems, recent studies have demonstrated the feasibility of using next generation sequencing (NGS) data including transcriptomics (e.g., RNA-seq ^18^) and proteomics ^19^ for categorizing MB and developed in-silico predictive methods to facilitate a more efficient and convenient prediction process. Previous studies indicate that identifying MB subtypes based on transcriptomic data is an efficient and feasible way ^20^ as it is an essential tool for expression profiling and provides a way for comprehensive understanding of biological problems ^21^. Most recently, medulloPackage ^22^ is constructed for rapid and accurate MB subtype identification based on the RNA-Seq and microarray datasets. medulloPackage identified discriminating features through gene ratio analysis and an unweighted mean of normalized scores is employed for sample classification. However, this approach would introduce bias as the data did not necessarily conform to normal distribution. In addition, another method named Medullo-Model ToSubtype (MM2S) ^23^ is developed to enable classification of individual gene expression profiles from MB samples, including patient samples, mouse models, and cell lines, against well-established molecular subtypes. MM2S achieves high accuracy for well-characterized subtypes like WNT and SHH. However, its performance is limited for Group 3 and Group 4, which remain poorly characterized and more heterogeneous, leading to challenges in accurate classification. In summary, existing MB subtyping methods directly use absolute gene expression levels without accounting for batch effects across different cohorts, which may lead to inaccurate classifications. Although transcriptomics datasets with varying compositions and sample sizes have been made publicly available, effectively utilizing diverse data sources remains challenging due to the severe batch effects. Various methods for batch effect adjustment, such as SVA ^24^, svaseq ^25^, DWD ^26^, XPN ^27^, and RUV ^28^, have been proposed, while normalizing across different batches can still distort true biological signals, particularly when sample distribution between batches is uneven ^29–32^.

Instead of relying on absolute gene expression levels, strategies like pair-wise analysis of gene expression (PAGE) ^33^ leverages the gene expression ratio (GER) ^34^ which exhibits consistent patterns within samples to construct the model ^35^ for batch effect correction ^36,37^. The GER strategy has been widely utilized in various biomedical domains ^35,38,39^, as its stability allows GER to be applied across different technological platforms, enhancing its utility in comparative omics studies. For instance, Zhao et al. ^37^ developed DRGpair which based on intra-sample paired GER for ovarian cancer detection to overcome the limitations of absolute gene expression values including batch effects and biological heterogeneity. Another transfer learning method, scPAGE2 ^36^, based on intra-sample GER using PAGE strategy to enhance generalization for sepsis diagnosis. By converting discrete gene pair differences into continuous GER values, scPAGE2 integrates both single-cell and bulk RNA-seq data. Moreover, recent studies ^38–40^ suggest that ranking gene expression, unlike absolute values which are sensitive to batch effects, provided a more stable representation of gene expression patterns. Additionally, ranking is scale-independent and it proceeds multi-cohort integration regardless of the difference among profile measurements.

Herein, we propose a novel and accurate computational approach called RaMBat for MB subtyping across diverse transcriptomics datasets with severe batch effects, based on intra-sample gene expression ranking information as well as a series of steps for gene-rank based information processing, enabling effective batch effect correction and robust integration of multiple bulk transcriptomics datasets. Based on 13 heterogeneous transcriptomics datasets, we demonstrate the accuracy and efficiency of RaMBat by comparing it with state-of-the-art methods like medulloPackage ^22^ and MM2S ^23^ as well as other machine learning (ML) methods equipped with the conventional batch correction method ComBat ^41^. Additionally, we have developed an R package RaMBat that is available on GitHub at https://github.com/wan-mlab/RaMBat.

## Results

### Study design for MB identification

To get subtype-specific signatures for MB samples across different cohorts with severe batch effects, we proposed RaMBat to identify MB subtype accurately from diverse transcriptomics data sources based on the gene expression ranking. The overall workflow contained four key steps (**Fig. 1)** which were designed to efficiently capture both global and local dependencies within high-dimensional intra-sample gene expression data from different cohorts, including (1) intra-sample gene expression analysis, (2) reversed expression pattern analysis, (3) informative GERs selection and (4) MB sample subtyping. As input, transcriptomics data was firstly converted into gene expression ranking followed with intra-sample gene expression analysis which was a two-step process including gene rank analysis and two-sided t-test (See Methods; **Fig. 1A**). Intra-sample gene expression analysis was performed between each pair of subgroups to identify genes that were specifically and significantly associated with each subtype. Next, the reversed expression pattern analysis (See Methods; **Fig. 1B**) was performed as a two-step process including reversal ratio analysis and Fisher’s exact test. Reversal ratio analysis leveraged rank levels between every possible pair of genes to retrieve subtype-specific GERs, combining each two genes together to construct the GER matrix. GER contained in each subtype was compared exhaustively to find the reversed gene expression pattern between each two subtypes cross samples. To enhance the accuracy for MB subtyping, LASSO ^42^ (see Methods; **Fig. 1C**) was applied to identify non-zero coefficients GERs that were strongly associated with subgroups. Similarly, as described above, comparisons between each pair of subgroups were performed and the identified GERs formed the foundation of RaMBat.

**Fig. 1.**
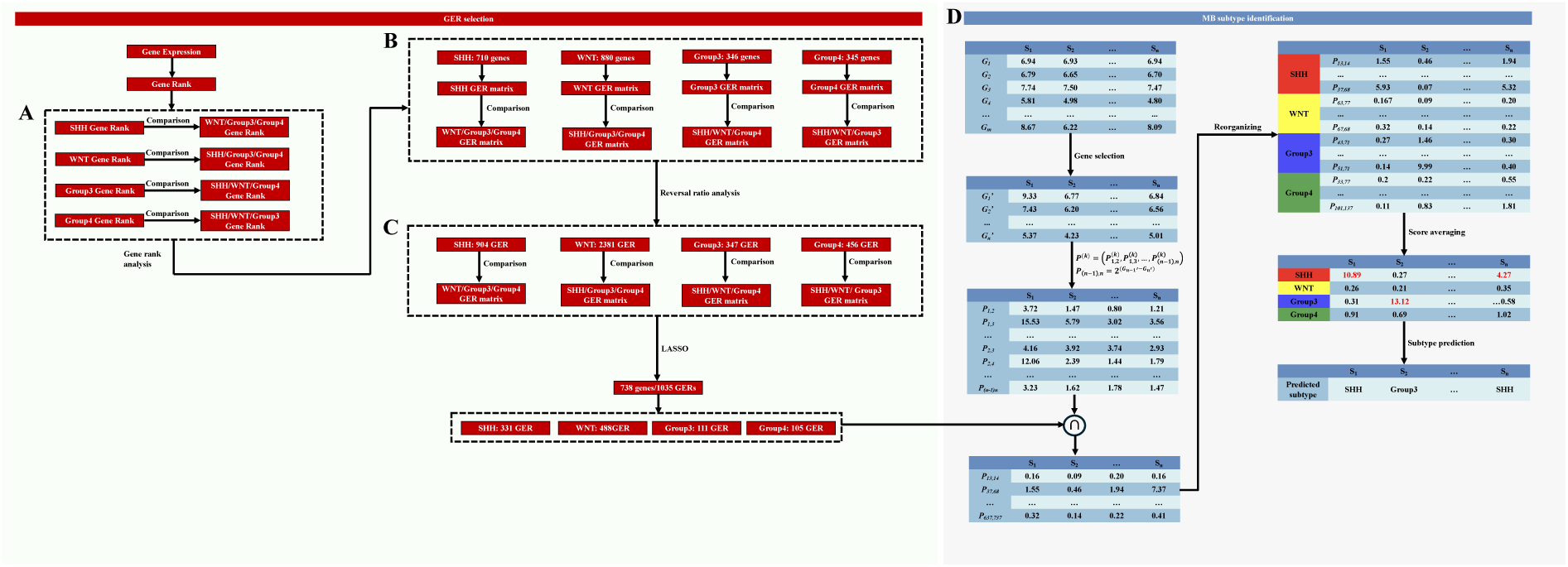
The workflow of RaMBat based on GERs for MB subtyping. RaMBat consisted of four steps for GERs selection and MB sample subtyping. (**A**) Perform gene ranking analysis. (**B**) Calculate reversal ratio values for GERs filtering. (**C**) Select GERs using LASSO. (**D**) Identify MB subtypes based on selected GERs.

In this study, a large-scale microarray dataset (GSE85217, 763 samples) was used to training. To ensure robust performance evaluation, RaMBat was tested on 13 independent test datasets which were generated using different platforms and exhibited significant scale disparities, necessitating standardized normalization to enhance comparability. 1035 GERs, corresponding to 738 unique genes (**Fig. 1A-C**), were identified. WNT had the highest number of GERs, totaling 488, followed by 331, 111, 105 GERs for SHH, Group 3 and Group 4, respectively. Subtype information was determined by calculating the average GER score for each subtype within each sample, with the subtype corresponding to the highest average GER score being assigned as the subtype for the sample (see Methods; **Fig. 1D**).

### Benchmarking with the state-of-the-art methods for identifying MB subtypes

To demonstrate the superiority of RaMBat, we benchmarked RaMBat with state-of-the-art methods including medulloPackage and MM2S as well as seven ML classifiers for MB subtyping across 13 independent datasets. Among them, medulloPackage was developed based on RNA-seq and microarray datasets for MB subtype prediction and MM2S was constructed based on microarray data for classification of patient samples, mouse models, and cell lines. Additionally, we trained and optimized seven ML classifiers, including SVM, LR, RF, XGBoost, KNN, NB and MLP, as benchmarks for comparison. In general, RaMBat consistently performed the best among all the benchmarking methods for MB subtyping. Specifically, RaMBat outperformed medulloPackage and MM2S in eight (**Fig. 2B-E, G, I, K, N**) of 13 datasets (**Fig. 2B-N**) and 11(**Fig. 2B-F, H, J-N**) out of 13 datasets respectively. For the remaining five (**Fig. 2F, H, J, L, M**) and two (**Fig. 2G, I**) datasets, respectively, RaMBat achieved the same accuracy as these methods. The overall accuracy for RaMBat across all datasets was 99.02%, where three Group 3 samples in GSE21140 and one Group 3 sample in GSE10327 were misclassified as Group 4, and one Group 4 sample in GSE74195 was misclassified as Group 3. As previously noted, the high similarity in transcriptomic and genomic profiles between Group 3 and Group 4 resulted in the molecular ambiguity and histological misclassification. However, the overall accuracy was still 11.8% (absolute) higher than MM2S and 7.9% (absolute) higher than medulloPackage. Additionally, the overall subtyping performance of RaMBat was better than all seven ML classifiers (**Fig. 2A**). The lowest accuracy achieved by RaMBat was 95% in GSE74195 (**Fig. 2J**), and the highest accuracy of 100% was recorded in ten datasets, including GSE73038 (**Fig. 2B**), GSE67850 (**Fig. 2E**), GSE50765 (**Fig. 2F**), GSE49243 (**Fig. 2G**), GSE30074 (**Fig. 2H**), GSE37382 (**Fig. 2I**), GSE62803 (**Fig. 2K**), GSE41842 (**Fig. 2L**), GSE50161 (**Fig. 2M**), GSE12992 (**Fig. 2N**). When evaluating performance by subtype, RaMBat always outperformed medulloPackage and MM2S on all four subtypes across all independent datasets in terms of accuracy, precision, sensitivity, F1-score, MCC, G-measure, AUC, specificity, and Jaccard index (**Fig. 3**).

**Fig. 2.**
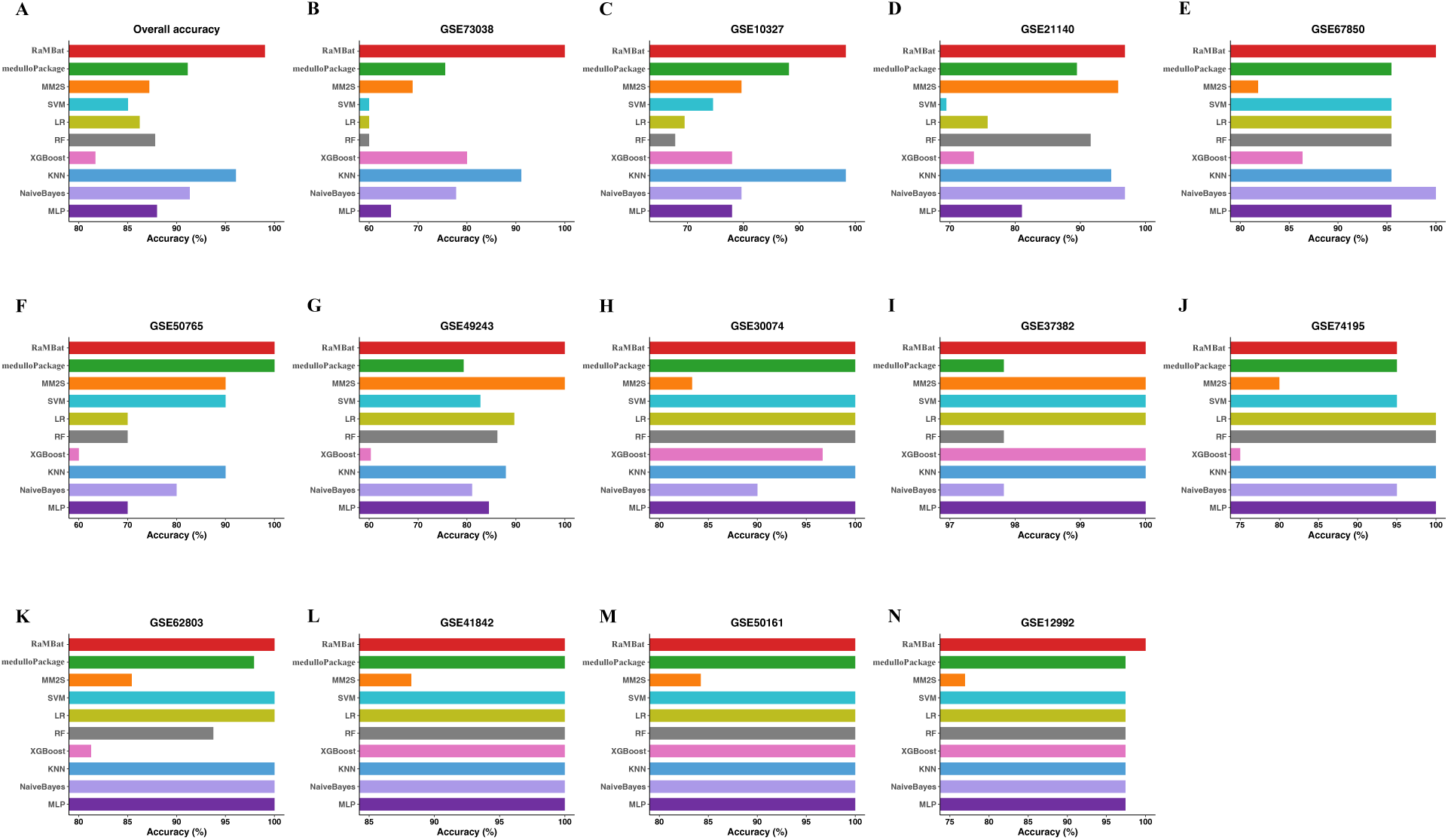
RaMBat significantly outperformed the state-of-the-art methods for MB subtyping based on 13 independent test datasets with severe batch effects. (**A**) Overall accuracy of RaMBat, medulloPackage, MM2S and other 7 ML classifiers were compared across all 13 independent datasets. Comparative performance of RaMBat, medulloPackage, MM2S and other 7 ML classifiers based on (**B**) GSE73038, (**C**) GSE10327, (**D**) GSE21140, (**E**) GSE67850, (**F**) GSE50765, (**G**) GSE49243, (**H**) GSE30074, (**I**) GSE37382, (**J**) GSE74195, (**K**) GSE62803, (**L**) GSE41842, (**M**) GSE50161, (**N**) GSE12992 in terms of accuracy, respectively. MLP, multilayer perceptron; KNN, k-nearest neighbors; XGBoost, eXtreme gradient boosting; RF, random forest; LR, logistic regression; SVM, support vector machine; MM2S, medullo-model to subtypes.

**Fig. 3.**
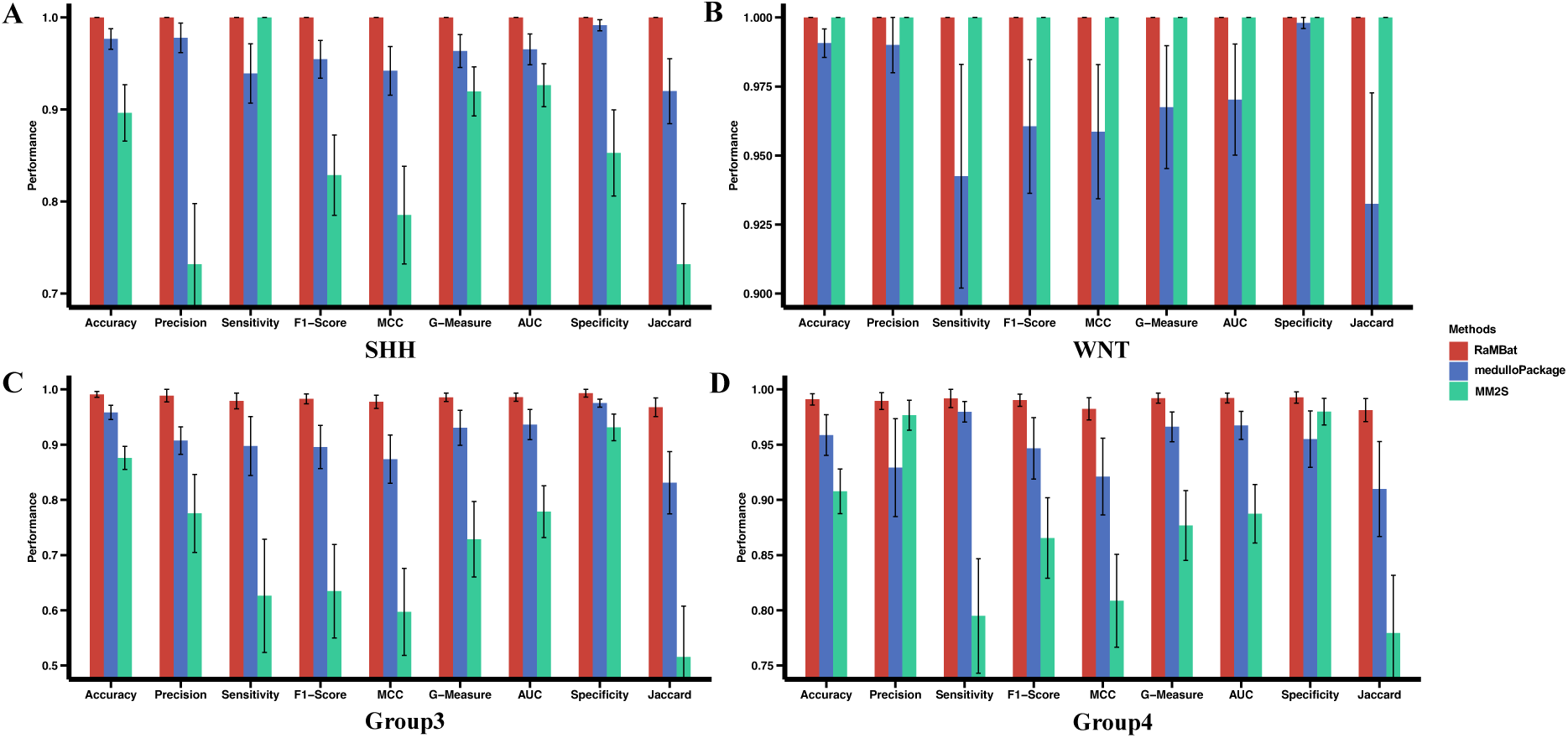
RaMBat achieved superior subtype-specific performance compared to state-of-the-art methods across 13 independent datasets for MB subtyping. Comparing performance of RaMBat with medulloPackage and MM2S across datasets in terms of accuracy, precision, sensitivity, F1-Score, MCC, G-measure, AUC, specificity and Jaccard index for SHH (**A**), WNT (**B**), Group 3 (**C**) and Group 4 (**D**). Metrics were obtained based on the prediction result of 508 samples in all 13 independent test datasets. MCC, Matthews correlation coefficient; AUC, area under the curve; Jaccard, Jaccard index

To further evaluate the impact of batch effects and the ability of RaMBat to efficiently integrate multi-source data with severe batch effects for MB subtyping, we combined all 13 independent test datasets and visualized the sample distribution using t-SNE (**Fig. 4**). When applied to raw gene expression data, conventional t-SNE visualization exhibited severe batch effects, with samples clustering predominantly by datasets rather than by biological subtypes (**Fig. 4A**). However, these technical variations obscured the biologically meaningful clustering of MB subtypes as the data did not cluster together according to major MB subtypes (**Fig. 4B**). In contrast, RaMBat effectively removed batch effects, enabling a clear separation of MB subtypes across diverse datasets (**Fig. 4C-D**). After integrating predicted subtype information from RaMBat into visualization, samples were well-integrated across different datasets (**Fig. 4C**), indicating that the batch effects were successfully removed. More importantly, RaMBat presented a clear visualization global landscape of all MB samples with all major subtypes distinctly separable (**Fig. 4D**). These results indicated that RaMBat can accurately integrate diverse bulk transcriptomics datasets, effectively removing batch effects while maintaining the underlying biological signals crucial for MB subtype classification.

**Fig. 4.**
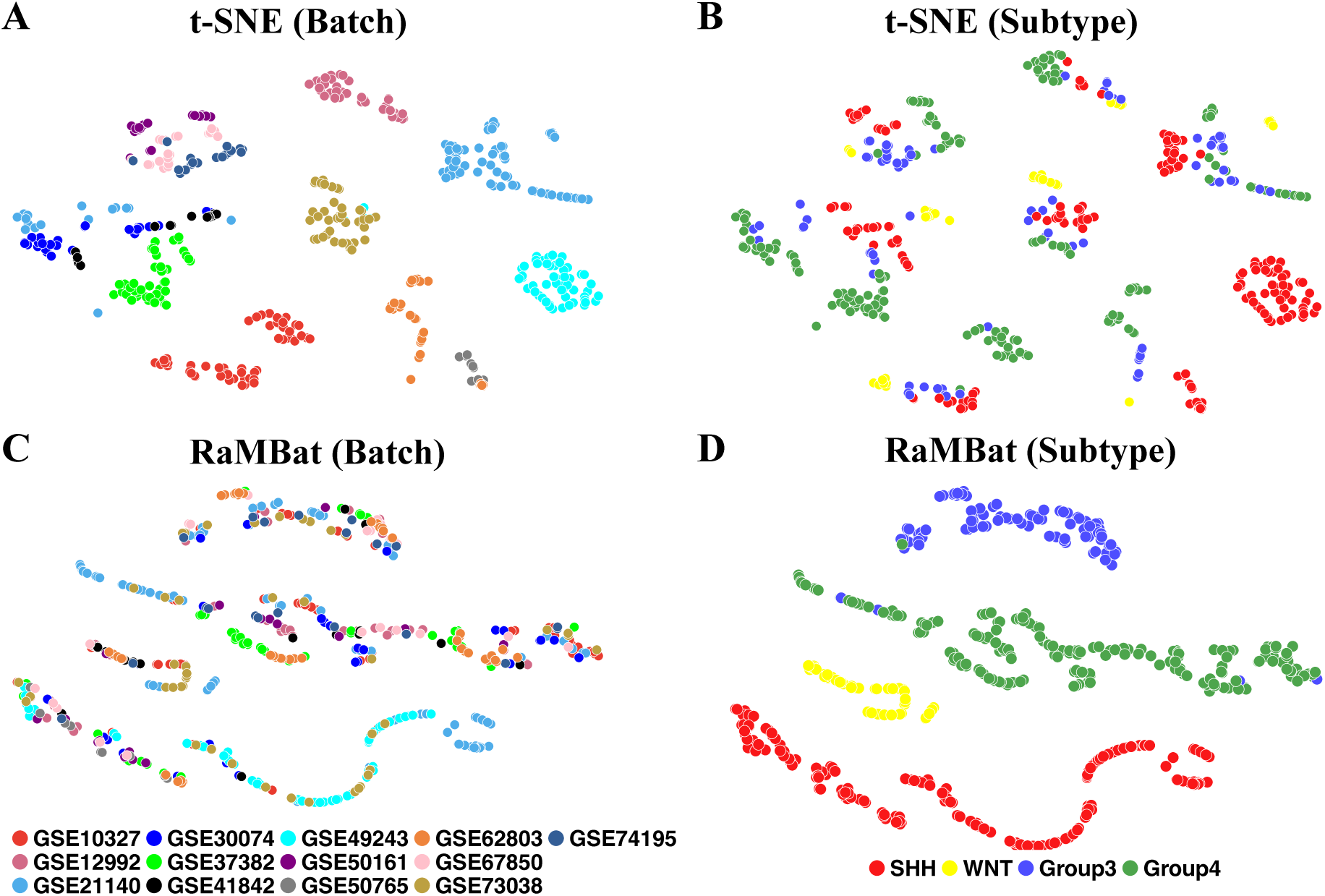
RaMBat enabled more effective visualization of MB subtypes than conventional t-SNE across 13 batch-affected independent datasets. (**A**) Samples were labeled by batch based on all samples in independent test datasets without normalization. (**B**) Samples were labeled by subtype based on all samples in independent test datasets without normalization. (**C**) RaMBat visualization in which samples were labeled by batch based on all samples in normalized independent test datasets. (**D**) RaMBat visualization in which samples were labeld by subtype based on all samples in normalized independent test datasets.

### Subtype-specific GERs selection

To investigate whether the marker genes identified by RaMBat were biologically meaningful, we used the training dataset as the basis for GER selection. The absolute expression levels were converted into intra-sample rankings, followed by gene rank analysis and two-sided t-test. There were 1055 ranking down genes and 1257 ranking up genes between SHH and Group 3, 1123 ranking down genes and 1233 ranking up genes between SHH and Group 4, 1220 ranking down genes and 1201 ranking up genes between SHH and WNT, 1221 ranking down genes and 1325 ranking up genes between WNT and Group 3, 1476 ranking down genes and 1522 ranking up genes between WNT and Group 4, 632 ranking down genes and 561 ranking up genes between Group 3 and Group 4 (**Supplementary Fig. S1A-F**). After intersecting these ranked up and down genes, 710, 880, 346, and 345 differentially ranked genes were identified for the SHH, WNT, Group 3, and Group 4, respectively.

Based on the differentially ranked genes in each subtype, 251,695, 386,760, 59,685 and 59,340 GERs for SHH, WNT, Group 3 and Group 4 were constructed, respectively. For each subtype, we performed the reversed expression pattern analysis between each pair of subgroups. For each comparison, a contingency table was constructed for the Fisher’s exact test. A total of 904, 2,381, 347 and 456 GERs (**Fig. 1C**) were screened for SHH, WNT, Group 3 and Group 4 respectively. Based on the reversed GERs, LASSO feature selection was performed to select the most important features which results in a final list of 1,035 GERs, corresponding to 738 unique genes (**Fig. 1C**). To comprehensively analyze the subtype-specific gene expression landscape in MB, the gene frequency across all 1,035 GERs for each subtype was also analyzed (**Fig. 5A-D**). Specifically, the top 10 most frequent genes in each subtype were labeled. For example, potential marker genes identified for SHH include *NEUROG1* and *ATOH1* ^43^, while *FZD10* was highlighted as a potential marker for the WNT subtype ^44^. To further investigate the gene expression patterns underlying MB subtype differentiation, differential expressed gene (DEG) analysis was performed and DEGs for each subtype were identified (**Fig. 5E-H**). To assess the overlap between RaMBat selected genes and DEGs, the identified DEGs with 738 unique genes were intersected (**Fig. 5I-P**). Notably, all top 10 most frequently ranked genes (**Fig. 5A-D**) were present within the intersected genes, except for *ZDHHC15* in Group 3, demonstrating strong consistency between RaMBat’s feature selection and conventional DEG analysis. Notably, RaMBat identified important genes that were not captured by conventional DEG analysis, such as *BOC*, which has been implicated in promoting the progression of early MB to more advanced stage. This finding suggested that RaMBat effectively captured subtype-specific molecular signatures, boosting its reliability in MB classification. To further validate the robustness of these findings, the expression patterns of the top 10 intersected genes of each subtype were tested using heatmaps. Distinct expression patterns of these 738 differentially ranked genes (**Fig. 6A)** evidenced their ability to effectively distinguish each MB subtype from others and enable clear separation into specific subgroups. The hierarchical clustering results further confirmed the biological relevance of the selected genes in MB subtyping. Specifically, based on data from GSE85217, the expression profiles of the top 10 most frequently ranked intersected genes were distinct across MB subtypes (**Fig. 6B-E)**. The results indicated specific expression patterns that effectively distinguished MB subtypes. Additionally, the expression level of upregulated DEGs, including *PDLIM3*, *TMEM132C*, *GABRA5* and *RBM24*, across all 13 independent test datasets highlighted their specific overexpression in SHH, WNT, Group 3 and Group 4 respectively (**Fig. 6F-I).**

**Fig. 5.**
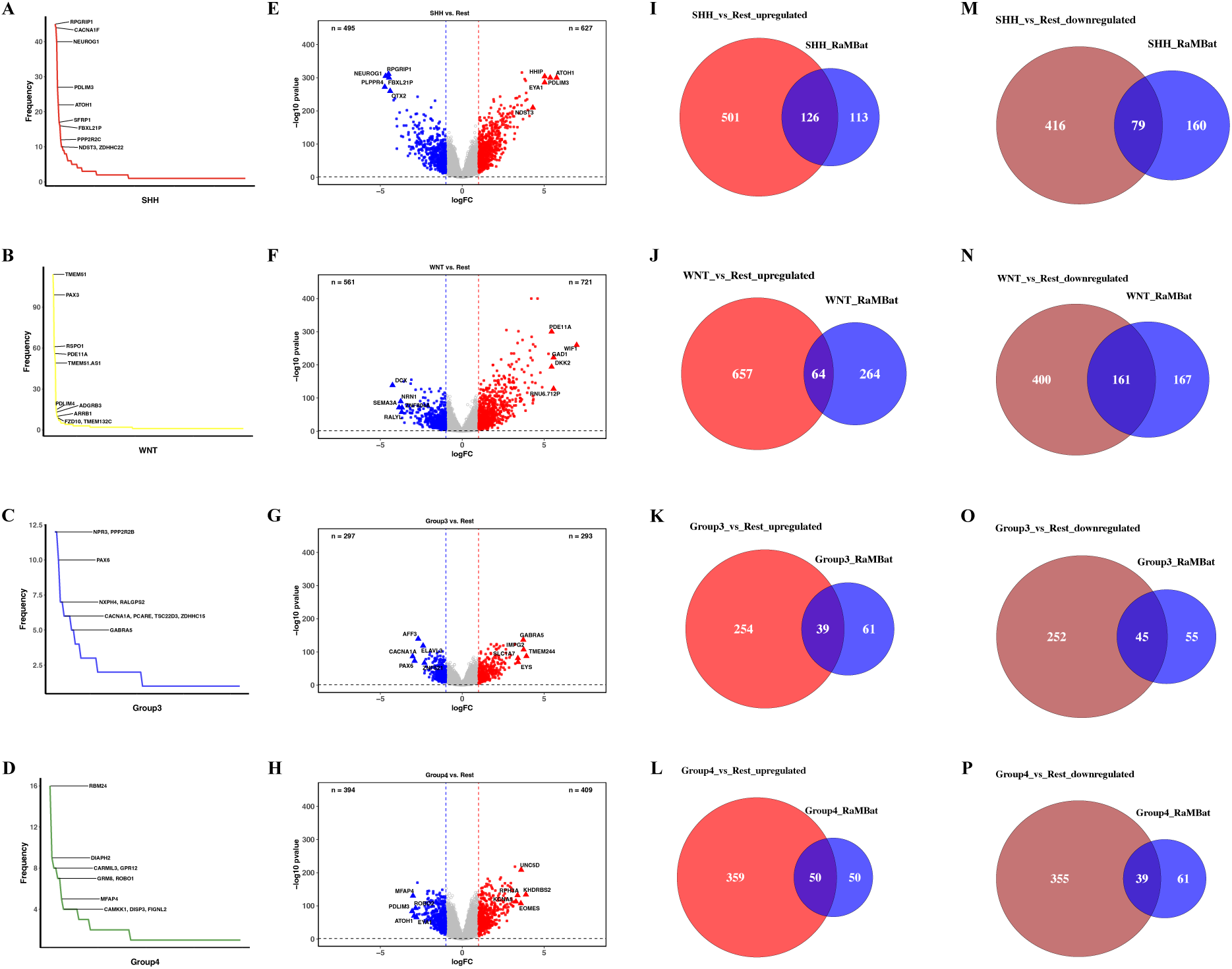
MB subtype-specific differential expression analysis showed strong concordance between RaMBat-selected features and conventional DEG analysis. The frequency distribution of all unique genes included in GERs for SHH (**A**), WNT (**B**), Group 3 (**C**), and Group 4 (**D**) respectively. The top 10 most frequently ranked genes in SHH, WNT, Group 3, and Group 4 were labeled. Differential expressed genes for SHH (**E**), WNT (**F**), Group 3 (**G**), and Group 4 (**H**) were identified. The x-axis represented the log2 fold change while y-axis showed -log10(p-value). Red and blue dots indicated significantly upregulated and downregulated genes respectively. Triangles highlighted top 5 upregulated and downregulated genes in each comparison. (**I-L**) Genes consistently identified by both DEG analysis and RaMBat. The intersections represented genes identified by both DEG analysis and RaMBat for SHH (**I, M**), WNT (**J, N**), Group 3 (**K, O**), Group 4 (**L, P**), where panels **I-L** showed upregulated genes, and panels **M-P** showed downregulated genes.

**Fig. 6.**
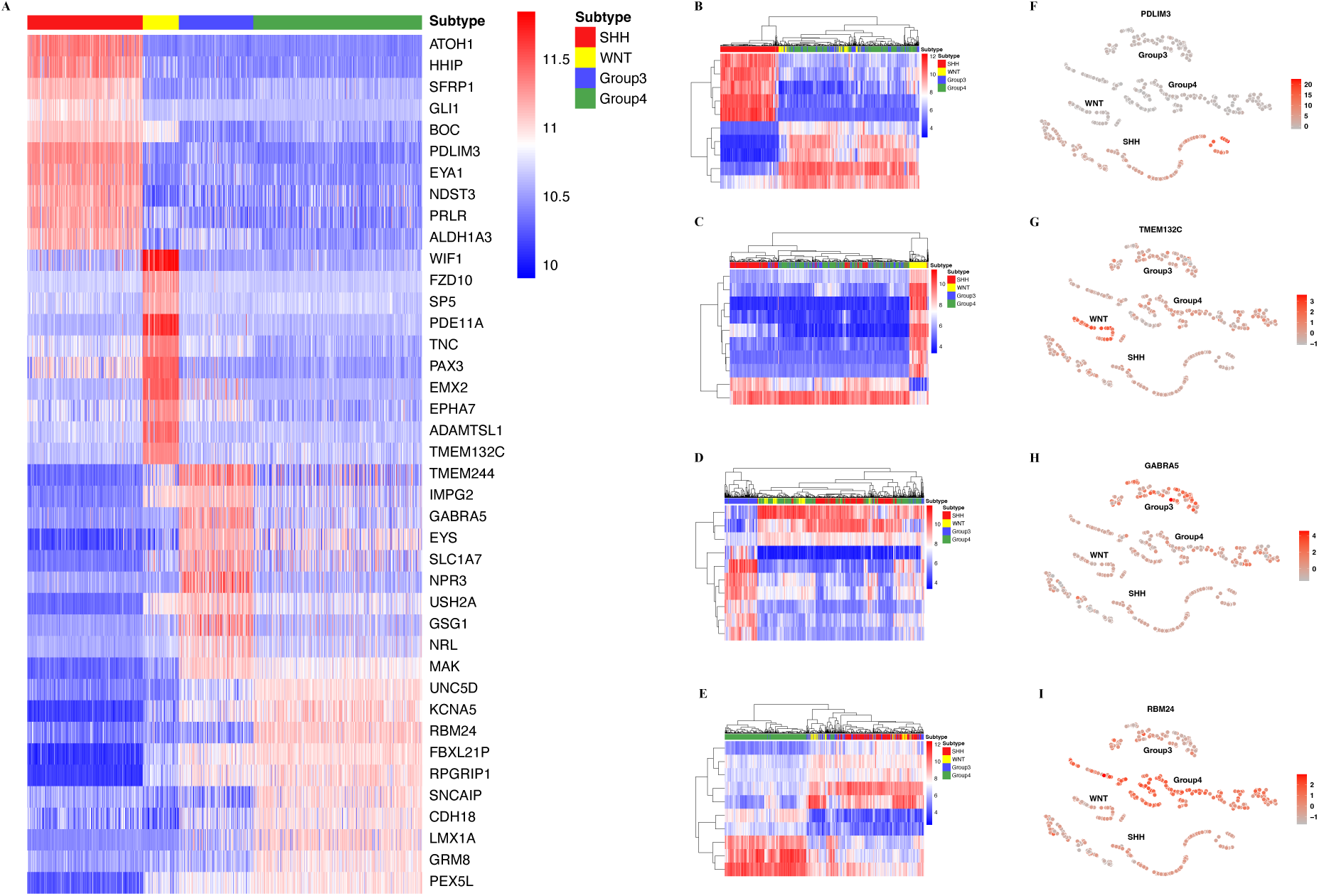
Identified differentially ranked genes showed distinct subtype-specific gene expression patterns. (**A**) The expression pattern of 738 unique genes associated with 4 molecular subtypes across samples in GSE85217. The absolute gene expression levels of the top 10 frequently ranked genes for SHH (**B**), WNT (**C**), Group 3 (**D**), and Group 4 (**E**) respectively. For all heatmaps, rows corresponded to genes, and columns corresponded to samples in GSE85217. Color scale ranged from blue (low expression) to red (high expression). (**F-I**) The expression level of the top upregulated DEG for SHH (**F**), WNT (**G**), Group 3 (**H**), and Group 4 (**I**) across 13 independent datasets with normalization. Each point represented a sample, colored by expression intensity (red: high, grey: low).

## Discussion

Medulloblastoma is a highly malignant pediatric brain cancer in cerebellum with peak incidence between ages 3-7 and a slight male predominance ^3^. Based on the molecular characteristics, it is mainly classified into four subtypes: SHH, WNT, Group 3 and Group 4, each with specific genetic profiles and prognostic outcomes. These four subtypes have varied symptoms to each other and exhibit significant variations in treatment response and prognosis, therefore, classifying patients into accurate subtype plays significant roles in the following therapy. Computational methods are designed to efficiently predict accurate subtype information. However, the absolute gene expression level deviated due to batch effect and biological heterogeneity even if using appropriate normalization methods. Previous studies ^45,46^, like iPAGE ^33^, has evidenced the analysis of the intra-sample relative expression level between genes is robust and generalizable in detecting biological signals. Furthermore, gene pair-based analysis is able to perform cross-platform comparison by integrating absolute expression from different cohorts and consequently stable across experimental assays and platforms ^33,47^. Building upon these advantages, we develop RaMBat, which achieves better prediction performance than existing state-of-the-art methods for MB subtyping. The benchmark comparison among RaMBat, medulloPackage, MM2S and other optimized ML classifiers demonstrates that analyzing differentially ranked genes provides robust and reliable insights for MB subtyping, complementing traditional differential expression-based approaches.

RaMBat presents high accuracy across 13 independent test datasets, with the most misclassifications occurring between Group 3 and Group 4. Among all test samples, one Group 4 sample is misclassified as Group 3, and four Group 3 samples are misclassified as Group 4. Previous studies have noted that these two subtypes exhibit highly similarity in transcriptomic and genomic profiles, contributing to the ambiguity between these two subtypes. Although recent studies have uncovered the heterogeneity within Group 3 and Group 4 and even suggesting their division into eight subtypes ^16^, patients with Group 3 or Group 4 are currently still treated with same therapeutic regimen ^1,3^. However, there are ongoing studies attempting to stratify treatment by molecular subtype to improve outcomes.

RaMBat effectively identifies subtype-specific marker genes, demonstrating its robustness in selecting biologically relevant features for MB subtyping. Specifically, selected GERs contain *NEUROG1, GLI1, ATOH1* associated with the SHH signature. *NEUROG1* and *ATOH1* are the top-selected genes with 40 and 22 GERs respectively, whereas *GLI1* is only involved in one GER. This aligns with the study published in 2007 which indicates *ATOH1, NEUROG1* and *GLI1* might be relevant biomarkers for SHH subtype ^43^. *GLI1* is a transcription factor and a key downstream effector of SHH signaling, directly linking it to SHH-driven tumorigenesis. *ATOH1* is a well-established marker of cerebellar granule precursor cells, and its expression is strongly associated with medulloblastomas that arise from these cells. In contrast, *NEUROG1* highly expressed tumors are suggested to originate from cerebellar ventricular zone progenitors rather than granule cell precursors. Additionally, canonical over-expressed genes *HHIP, BOC* and *SFRP1* are also included in the GERs ^48^. The WNT subtype of medulloblastoma is closely linked to the activation of the WNT signaling pathway, primarily through mutations that stabilize beta-catenin and drive oncogenesis. Five WNT signaling genes are observed within GERs associated with the WNT subtype, including *SP5, WIF1, FZD10, CER1,* and *PRKCB*, which play various roles in the WNT signaling pathway. For example, the activation of the WNT pathway leads to the upregulation of *SP5*, which in turn promotes the expression of genes that contribute to tumorigenesis ^49,50^. Its presence indicates active WNT signaling, which is characteristic of the WNT subtype. The *WIF1* plays a significant role in medulloblastoma by acting as a negative regulator of the WNT signaling pathway ^51^. As a receptor in the Frizzled family, *FZD10* mediates WNT ligand binding, activating the canonical WNT/β-catenin pathway ^44^. Although less is known about the specific drivers, pathways, and biomarkers that discriminate Group 3 and Group 4 subtypes, it is established that these tumors share certain common pathways. For example, *UNC5D* has been identified as a key gene associated with neuronal migration in MB, showing specificity for certain subtypes and being present in GERs ^51,52^. Additionally, specific chromosomal aberrations affecting *AFF3* may contribute to tumorigenesis by disrupting regulatory pathways. The *AFF3* gene can potentially discriminate these two subtypes, as it is downregulated in Group 3, but upregulated in Group 4. As expected, it only presents in GERs for Group 3 ^51^. Thus, the detailed function of *AFF3* may worth more investigation.

Unlike absolute gene expression levels which are sensitive to batch effects, RaMBat is scale-independent and multi-cohort integration regardless of the difference among profile measurements. The data visualization by RaMBat (**Fig. 4**) further confirmed that it can efficiently eliminate batch effects and clearly separate MB subtypes, unlike conventional approaches such as t-SNE, which are susceptible to batch-induced variations. We admit that one limitation of our study is its high computational cost and time-consuming nature, particularly when handling large-scale datasets, which significantly reduces efficiency. Overall, this study proposes RaMBat, a novel approach leveraging ranking gene expression to select robust and subtype-specific GERs for accurate MB subtyping across diverse transcriptomics datasets with severe batch effects. Benchmarking across 13 datasets demonstrates that RaMBat outperforms state-of-the-art MB subtyping methods, achieving a high median accuracy of 99.02%, while also providing effective batch effect correction. We believe that RaMBat will significantly enhance downstream risk stratification for MB and contribute to the design of more personalized treatment strategies. Additionally, we also expect that RaMBat will be highly effective in addressing many biomedical problems with batch effects for various types of pediatric, adolescent, and young adult (AYA) cancers.

## Methods

### Datasets

A transcriptomic dataset (GSE85217) was used to train the RaMBat, characterized using Affymetrix Human Gene 1.1 ST Array and downloaded from the Gene Expression Omnibus ^53^ database (GEO, https://www.ncbi.nlm.nih.gov/geo). For model performance evaluation, 13 datasets were downloaded through the *getGEO* function in R with following accession numbers: GSE10327, GSE12992, GSE21140, GSE30074, GSE37382, GSE41842, GSE49243, GSE50161, GSE50765, GSE62803, GSE67850, GSE73038, GSE74195. Specifically, to address scale disparities, datasets were normalized as (*x* − *x̄*)/*s*, where each gene expression value *x* was subtracted by the mean value *x̄* of the column and dividing it by the standard deviation *s* of the column. In both the training and independent test datasets, Group 4 samples represented the largest proportion, followed by SHH, Group 3, and WNT. Additionally, the number of samples for each subtype was consistently higher in the training dataset compared to the independent test dataset. The training dataset, GSE85217, included 763 MB samples (223 SHH, 70 WNT, 144 Group 3, and 326 Group 4), while the independent test datasets contained 508 samples (177 SHH, 42 WNT, 92 Group 3, and 192 Group 4) (**Supplementary Table 1**).

To identify MB subtypes based on multiple bulk transcriptomics data, an accurate MB subtyping approach based on gene ranking for heterogeneous data with severe batch effects, named RaMBat, was developed. RaMBat contained four key steps (**Fig. 1A**), including intra-sample gene expression analysis, reversed expression pattern analysis, informative GERs selection, and finally MB sample subtyping. It leveraged gene expression rankings instead of absolute gene expression levels to address batch effects from different data sources. Based on 13 transcriptomics datasets with severe batch effects, we demonstrated that RaMBat significantly outperformed state-of-the-art methods like medulloPackage and MM2S as well as seven optimized ML classifiers equipped with the conventional batch correction method ComBat in terms of prediction accuracy for identifying MB subtypes.

### Intra-sample gene expression analysis

Based on the ascending order, from the smallest to the largest values, we ordered all genes inside the dataset. Subsequently, we transformed the absolute gene expression levels

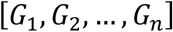

into rank values,

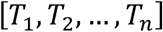

where *G* was the absolute value of gene expression, *n* was the total number of genes, *T* was the rank value of gene within one sample, where ranking was performed in ascending order. The gene with the lowest expression value was assigned a rank of 1, while the gene with the highest expression value was assigned a rank of *n*. We then divided the rank matrix of GSE85217 into four submatrices corresponding to the SHH, WNT, Group 3, and Group 4, based on sample labels. For each gene, we calculated the gene rank difference and a *p* value between each combination of submatrices, resulting in six different comparisons: SHH vs. WNT, SHH vs. Group 3, SHH vs. Group 4, WNT vs. Group 3, WNT vs. Group 4, and Group 3 vs. Group 4. The mean rank of each gene across all samples within one subtype was calculated and compared to its mean rank in another subtype, with the gene rank representing the difference between these two mean rank scores. A two-sided t-test was then performed to assess the statistical significance of the difference between subtypes. We applied thresholds of *p* < 0.05 and a gene rank > 2314 (representing 11% of the total number of genes) to identify significant differentially ranked genes.

### Reversed expression pattern analysis

To construct gene pairs based on the differentially ranked genes, we followed iPAGE ^33^ protocols to construct GERs. Specifically, for each sample, the rank values of all possible gene pairs were compared, encoding their relationships as binary indicators, represented by

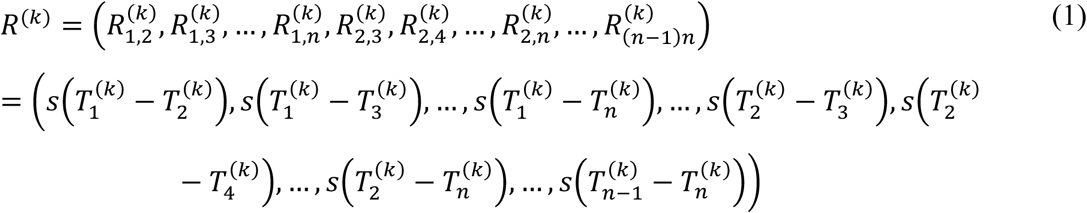

where GERs (*R*) was constructed by gene rankings (*T*) in a sample *k* and function *s* denoted the process of converting gene rankings into GER values as following:

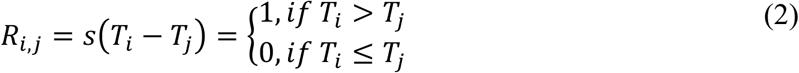

Every two different genes within each sample were subtracted to convert the rank matrix into GER matrix. Then, GERs with a significant difference in their relative expression *R*_*i,j*_ between each two subtypes were extracted.

Two situations were taken into consideration, where *R*_*i,j*_ = 1 and *R*_*i,j*_ = 0. We counted the number of samples with *R*_*i,j*_ = 1 and *R*_*i,j*_ = 0 in the first subtype group as *f*_1,1_ and *f*_1,2_respectively, then in the second subtype group, the number of samples with *R*_*i,j*_ = 1 and *R*_*i,j*_ = 0 as *f*_2,1_and *f*_2,2_respectively. Finally, the contingency table was shown as follows:

| | $T_i > T_j$ | $T_i \leq T_j$ |
| --- | --- | --- |
| Selected subtype | $f_{1,1}$ | $f_{1,2}$ |
| Rest subtype | $f_{2,1}$ | $f_{2,2}$ |

For example, for the comparison between SHH and WNT, *f*_1,1_ and *f*_1,2_ were the number of samples with *R*_*i,j*_ = 1 and *R*_*i,j*_ = 0 respectively in SHH, and *f*_2,1_ and *f*_2,2_ were the number of samples with *R*_*i,j*_ = 1 and *R*_*i,j*_ = 0 respectively in WNT.

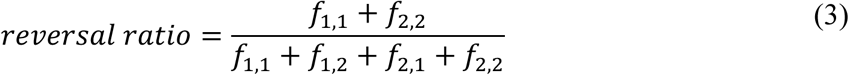

To calculate the reversal ratio, we considered 12 different comparisons, with each subtype having three pairwise comparisons. For example, when selected specific GERs for SHH, the comparisons were with WNT, Group 3, and Group 4, respectively. We established different thresholds for the reversal ratio based on the subtype comparisons. A threshold of 0.95 was set for the six pairwise comparisons within the SHH and WNT subtypes. For comparisons between Group 3 and SHH, Group 4 and SHH, we set the threshold at 0.9. A threshold of 0.85 was used for comparisons between Group 3 and WNT, Group 4 and WNT, Group 4 and Group 3. Lastly, a threshold of 0.8 was applied specifically for the comparison between Group 3 and Group 4. At the same time, we performed Fisher’s exact test to obtain the *p* value for each of the twelve comparisons. The *p* value was calculated by

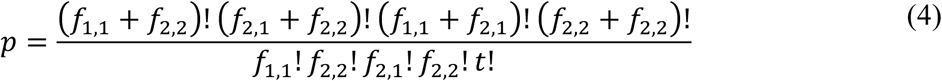

where *t* = *f*_1,1_ + *f*_1,2_ + *f*_2,1_ + *f*_2,2_ and the threshold for *p* value was set as 0.05. For each subtype, three sets of selected GERs were obtained which were then intersected to identify GERs consistently presenting across all three pairwise comparisons within that subtype. At the end of this step, subtype-specific GERs were selected for SHH, WNT, Group 3 and Group 4, respectively, based on a reversal ratio exceeding the threshold and a *p* value > 0.05 (**Fig. 1B**).

### Selecting informative GERs

To reduce the computational complexity of the subsequent MB sample subtyping step based on GERs and boost the stability of this method, we furthermore refined the number of GERs using LASSO (Least Absolute Shrinkage and Selection Operator) ^54^. LASSO naturally performed feature selection by shrinking the coefficients of less important features to zero, effectively eliminating them from the model. We conducted LASSO 12 times to select the most important GERs corresponding to three pairwise comparisons for each of the four subtypes (**Fig. 1C**). Similarly, we intersected three sets of selected GERs for each subtype. In the end, 1035 GERs, corresponding to 738 unique genes, with non-zero coefficients were filtered which were then used as the input for sample subtyping. LASSO was performed using R package “glmnet” ^55^. Additionally, based on these 1035 GERs, heatmap was plotted using the R package “pheatmap” ^56^.

### RaMBat Visualization

RaMBat utilized a combination of two key matrices: a subtype score matrix and a sample-to-subtype matrix derived from prediction results. The former matrix was obtained through the RaMBat subtype identification workflow (**Fig. 1**), in which GERs were used to calculate subtype-specific scores for each sample. This matrix captured the continuous prediction confidence of each sample across the four MB subtypes (SHH, WNT, Group 3, Group 4), forming a *R* × *C* matrix, where *R* is the number of samples and *C* is the number of subtypes. A sample-to-subtype matrix was constructed using one-hot encoding based on the predicted subtype labels derived from the subtype score matrix. Specifically, the subtype with the highest score for a given sample was assigned a value of 1, while all other subtypes were assigned 0. This results in a binary matrix of the same dimension (*R* × *C*), representing discrete subtype assignments. These two matrices were then concatenated along the feature axis to form a unified visualization matrix, which was used as input for t-SNE to generate a low-dimensional embedding. This integration strategy enabled the combination of both quantitative prediction certainty and categorical class identity, thereby enhancing the biological interpretability and subtype-level separation in the visualization space.

### Benchmarking with the state-of-the-art methods

To assess the prediction performance of RaMBat, we initially downloaded 13 independent test datasets with clear subtype information, containing 508 samples (**Supplementary Table 1**). The subtyping process for a given sample was composed of three stages, 1) convert the absolute gene expression into GERs based on the selected unique genes, 2) filter to the selected GERs in the model and calculate the mean score of GERs for each subtype, 3) determine the subtype with the highest score and assign it to the sample (**Fig. 1D**). Finally, we compared the performance of RaMBat with state-of-the-art MB classifiers, including the medulloPackage ^22^ and MM2S ^23,57^. medulloPackage was also developed using GERs but based on the differentially expressed genes (DEGs) selected by limma package in R. In contrast, MM2S employed a k-nearest neighbor (KNN) ^58^ classifier based on the gene-set enrichment analysis ranked matrix. Additionally, to compare the performance of RaMBat with machine learning methods, 7 classifiers, including support vector machine (SVM) ^59^, logistic regression (LR) ^60^, random forest (RF) ^61^, XGBoost ^62^, KNN, naïve bayes (NB), multilayer perceptron (MLP) ^54^ were trained and optimized. The training dataset and independent test datasets which were normalized and then batch effect corrected by the *combat* function in R were used for training and testing of these ML classifiers. For performance evaluation, we considered nine major metrics: accuracy, precision, sensitivity, specificity, area under the curve (AUC), Jaccard index (Jaccard), Matthews correlation coefficient (MCC), G-measure and F1-score.

## Supplementary Data

**Supplementary Fig. S1 Differential rank gene analysis within RaMBat.** (A-F) The differentially ranked gene for specific medulloblastoma subtypes. The x-axis represented the gene rank difference where y-axis showed -log10(p-value). Red dots indicated significantly up-ranked genes and blue dots represented significantly down-ranked genes. Triangles represented top 5 up/down-ranked genes. Differentially ranked genes between SHH and WNT (A), SHH and Group 3 (B), SHH and Group 4 (C), WNT and Group 3 (D), WNT and Group 4 (E), Group 3 and Group 4 (F).

**Supplementary Table 1 Comparing detailed information for each dataset**. Summary of MB subgroup distribution in training and independent test datasets. Num, number of samples.

## Author contributions

SW conceived and designed the study. SW and MS developed the method, performed the experiments and analyzed the data. All authors participated in writing the paper. SW and MS revised the paper. The manuscript was approved by all authors.

## Data availability

All the data used in this manuscript are publicly available in the corresponding references.

## Code availability

RaMBat is available as an R package in GitHub at https://github.com/wan-mlab/RaMBat.

