## supplementary files for "Accurate identification of medulloblastoma subtypes from diverse data sources with severe batch effects by RaMBat"

**Table 1 Comparing detailed information for each dataset.** Summary of MB subgroup distribution in training and independent test datasets. Num, number of samples.

| Cohorts | Platform | Num | | | | Total |
|---|---|---|---|---|---|---|
| | | SHH | WNT | Group3 | Group4 | |
| GSE85217 | Affymetrix Human Gene 1.1 ST Array | 223 | 70 | 144 | 326 | 763 |
| GSE10327 | Affymetrix Human Genome U133 Plus 2.0 Array | 14 | 9 | 10 | 26 | 59 |
| GSE12992 | Affymetrix Human Genome U133 Plus 2.0 Array | 7 | 4 | 8 | 20 | 39 |
| GSE21140 | Affymetrix Human Exon 1.0 ST Array | 29 | 8 | 23 | 35 | 95 |
| GSE30074 | Affymetrix Human Gene 1.0 ST Array | 9 | 2 | 3 | 16 | 30 |
| GSE37382 | Affymetrix Human Gene 1.1 ST Array | 10 | | 5 | 31 | 46 |
| GSE41842 | Affymetrix Human Gene 1.0 ST Array | 3 | 6 | 2 | 6 | 17 |
| GSE49243 | Affymetrix Human Genome U133 Plus 2.0 Array | 58 | - | - | - | 58 |
| GSE50161 | Affymetrix Human Genome U133 Plus 2.0 Array | 9 | 1 | 2 | 7 | 19 |
| GSE50765 | Affymetrix Human Gene 1.1 ST Array | 10 | - | - | - | 10 |
| GSE62803 | Affymetrix Human Gene 1.1 ST Array | 6 | 5 | 14 | 23 | 48 |
| GSE67850 | Affymetrix Human Genome U133 Plus 2.0 Array | 5 | 1 | 9 | 7 | 22 |
| GSE73038 | Affymetrix Human Genome U133 Plus 2.0 Array | 16 | 10 | 9 | 10 | 45 |
| GSE74195 | Affymetrix Human Genome U133 Plus 2.0 Array | 1 | 1 | 7 | 11 | 20 |

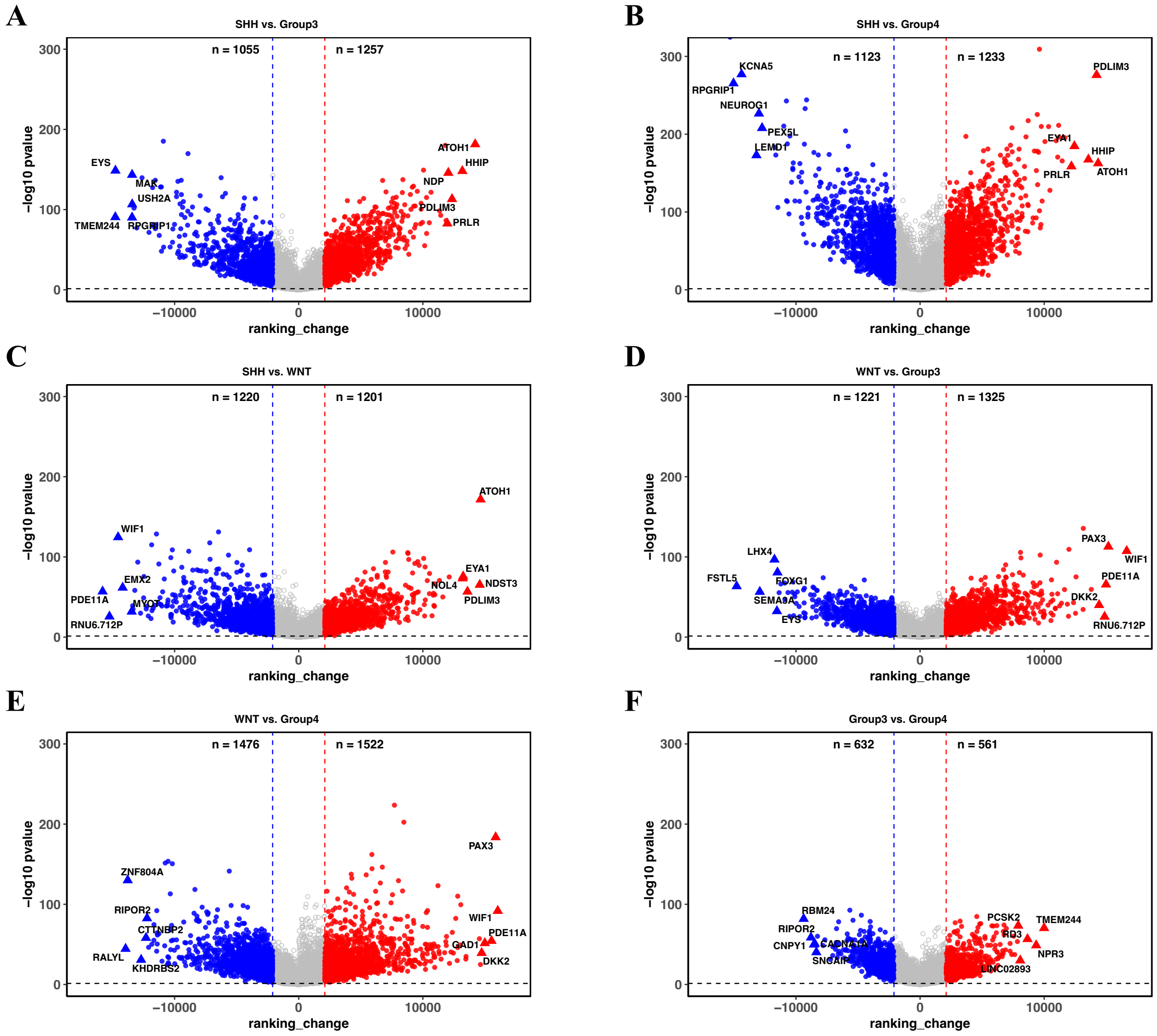


**Supplementary Fig. S1 Differential rank gene analysis within RaMBat.** (**A-F**) The differentially ranked gene for specific medulloblastoma subtypes. The x-axis represented the gene rank difference where y-axis showed -log10(p-value). Red dots indicated significantly up-ranked genes and blue dots represented significantly down-ranked genes. Triangles represented top 5 up/down-ranked genes. Differentially ranked genes between SHH and WNT (**A**), SHH and Group 3 (**B**), SHH and Group 4 (**C**), WNT and Group 3 (**D**), WNT and Group 4 (**E**), Group 3 and Group 4 (**F**).
